# Intergenerational paternal effect of adult density in *Drosophila melanogaster*

**DOI:** 10.1101/368613

**Authors:** Purbasha Dasgupta, Saubhik Sarkar, Akankshya A. Das, Tanya Verma, Bodhisatta Nandy

**Author notes:** Equal contribution (Purbasha Dasgupta, Saubhik Sarkar).

## Abstract

1. Notwithstanding recent evidences, paternal environment is thought to be a potential but unlikely source of fitness variation that can affect trait evolution. Here we studied intergenerational effects of males’ exposure to varying adult density in *Drosophila melanogaster* laboratory populations.
2. We held sires at normal (N), medium (M) and high (H) adult densities for two days before allowing them to mate with virgin females. This treatment did not introduce selection through differential mortality. Further, we randomly paired males and females and allowed a single round of mating between the sires and the dams. We then collected eggs from the dams and measured the egg size. Finally, we investigated the effect of the paternal treatment on juvenile and adult (male) fitness components.
3. We found a significant treatment effect on juvenile competitive ability where the progeny sired by the H-males had higher competitive ability. Since we did not find the treatment to affect egg size, this effect is unlikely to be mediated through variation in female provisioning.
4. Male fitness components were also found to have a significant treatment effect: M-sons had lower dry weight at eclosion, higher mating latency and lower competitive mating success.
5. While being the first study to show both adaptive and non-adaptive effect of the paternal density in *Drosophila*, our results highlight the importance of considering paternal environment as important source of fitness variation.

## Introduction

Parental environment has the potential to influence offspring traits and fitness through intergenerational effects (and more stable transgenerational effects, see Dias and Ressler 2014 for the distinction between trans and intergenerational effects). While it can potentially pass on deleterious effects of different components of the environment to the following generation (Yahuda et al. 2000), intergenerational effect can also be adaptive, especially under fluctuating environment (Bonduriansky and Day 2009). Among the myriad components of an organism’s ecology, few factors are as variable as density and nutritional availability. Both have been recently found to have intergenerational effects, especially through the maternal route (i.e., maternal effect) in a wide variety of organisms (Mousseau & Fox 1998). There is a growing body of evidence showing the importance of the intergenerational effect of paternal nutrition, social experience and density on fitness related traits of the offspring (Friberg et al., 2012; Adler & Bonduriansky, 2013; Crean et al., 2013, Dasgupta et al. 2016). However, the prevalence and adaptive significance of such paternal effect is yet to be ascertained.

There are many reports of environment dependent maternal effect mediated through variation in maternal provisioning in egg/offspring (Rossiter, 1996; Mousseau & Fox, 1998). For example, females living under high density may suffer from adverse effects of crowding (such as, malnutrition) and may therefore struggle to allocate resources in maternal provisioning either in the form of stored resources in egg or lactation, which in turn may lead to poor quality progeny (Christian & Lemunyan, 1958). Alternatively, females raised in high density may strategically produce fewer eggs/progeny while investing more resources (e.g., yolk) in each of them – thereby giving the progeny a better start for the impending challenges of crowding (Prasad et al., 2003; Holbrook & Schal, 2004; Mitchell & Read, 2005; Vijendravarma et al. 2010). Generally, under fluctuating environmental conditions, such parental ability to optimize offspring phenotype has been conjectured to be adaptive (Bonduriansky & Day, 2009; Kuijper & Hoyle, 2015). For example, Guppy (*Poecilia reticulata*) females were found to produce larger offspring (a) under food limitation (Reznick & Reznick, 1993) and (b) when they experienced high level of competition – priming the offspring for better competitive ability (Bashey, 2006). The larger eggs produced by *D. melanogaster* females that grew in nutritionally impoverished food, survive (egg to adult survivorship) better in impoverished food and give rise to smaller adults (Vijendravarma et al., 2010). In contrast, Valtonen et al. (2012) found *D. melanogaster* females grown on impoverished food to produce larger offspring (adult) compared to those grown on nutritionally rich food. Note that many of the maternal effects discussed above are mediated through variation in resource provisioning by mothers.

Not surprisingly, most of the reports of environment dependent paternal effect (intergenerational and transgenerational) come from animals with paternal provisioning through nuptial gift transfer to the females (Dussourd et al., 1988; Gwynne, 1988; Zeh & Smith, 1995; Smedly & Eisner, 1996; Vahed, 1998). However, it is only recently that studies have started to address if similar paternal effects are also present in species without paternal provisioning. In one of the first such explicit studies, female Neriid flies (*Teleostylinus angusticollis*) raised on richer diet were found to produce larger eggs and offspring that developed faster, while males raised on richer diet sired larger offspring with better survival rate, especially under resource scarcity (Bonduriansky & Head, 2007; Adler & Bonduriansky, 2013). In a solitary Ascidian, *Styela plecata*, males were found to produce offspring with phenotype corresponding to the population density experienced by the father (Crean et al., 2013). In fruit flies, *D. melanogaster*, Valtonen et al. (2012) reported that fathers fed on poor quality diet sire larger sons. Paternal experience of the intensity of competition (assessed by the number of co-inhabitant rival males) adaptively affected reproductive behaviour of male offspring in *D. melanogaster* (Dasgupta et al., 2016). Islam et al. (1994) showed paternal social environment to have a significant impact on offspring behavioural traits. Paternal experience of ambient temperature was also found to affect offspring fecundity in *D. melanogaster* (Huey et al., 1995). Low temperature was found to affect offspring phenotype in two other species of Drosophila – *D. simulans* (Watson & Hoffmann, 1995) and *D. serrata* (Magiafoglou & Hoffmann, 2003). Thus, there is a growing body of evidence showing environment dependent paternal effect. In addition to affecting viability, such paternal effect has been shown to affect progeny reproductive performance and hence is likely to be key player in sexual selection (for example, see Bondurianky & Head, 2007). However, such data are far from being plenty.

Here we investigated the effect of paternal experience of population density on progeny fitness components, including male mating behaviour in *D. melanogaster* laboratory adapted populations. As discussed previously, paternal effect has already been reported in these (Dasgupta et al., 2016) and other populations of *D. melanogaster*, establishing them as a relevant system to investigate the paternal effect and its consequences on Darwinian fitness (William et al., 2006). Further, laboratory adapted populations of *D. melanogaster* have been used to investigate the fitness consequence of a plethora of environmental parameters, including population density. Fruit flies naturally grow in ephemeral resource patches, such as rotting fruits and vegetables. Crowding in transiently available rich patches is expected to be a key component of their natural ecology. Density of adults in a resource patch not only determines the extent to which individuals must compete for food and limited space (e.g., oviposition substrate) but also for other resources, such as suitable mates. Increase in density also leads to an increase in the probability of disease transmission (Barnes & Siva, 2000). In essence, density often determines the nature and intensity of selection acting on a population and has been studied within the broader premises of density dependent selection (MacArthur & Wilson, 1967; Mueller, 1997; Prasad & Joshi, 2003). Much of the existing literature investigated adaptation to increased (but stable) juvenile or adult density using experimental evolution on laboratory populations of *D. melanogaster* (Mueller & Sweet, 1986; Mueller et al., 1991; Nagarajan et al., 2016; Sarangi et al., 2016; Shenoi et al., 2016; Shenoi & Prasad, 2016). However, little is known about adaptation to fluctuating density. Intergenerational and transgenerational effects, if used by the parents to optimize offspring phenotype, can be of adaptive value if density fluctuation across generation is, at least to some extent, predictable. Interestingly, these experimental evolution studies reported ‘rapid’ adaptation to ‘crowding’. Though evidences unequivocally showed the genetic changes associated with such adaptation, non-genetic parental effects (trans and intergenerational) may, in addition, account for the ‘rapid’ adaptation (Bonduriansky and Day 2009). However, this idea has not been tested – an existing lacuna in the literature, which we intend to fill to some extent.

To investigate the paternally transmitted intergenerational effect of varying density, we subjected males to three adult density treatments and then allowed them to sire progeny by mating the treated males to untreated dams. We then assessed the effect of the paternal adult density (hereafter, referred to as paternal density) treatment on progeny fitness components in juvenile (juvenile competitive fitness) and adult stages (males: mating ability, mating latency, copulation duration, courtship frequency, competitive mating success). We found the paternal density treatment to have significant intergenerational effect on both juvenile and adult fitness components.

## Materials and Methods

All the experiments were done using a set of laboratory adapted populations of *D. melanogaster* – BL. Full laboratory history of these populations can be found in (reference blinded). Briefly, these are a set of five replicate populations (BL _1-5_) maintained on standard Banana-Jaggery-Yeast food, under 14-day discrete generation cycle at 25 °C ambient temperature, 60-80% relative humidity, with population size ~2800. Larval density is maintained at ~70 per 6-8ml food per vial (25mm×90mm, diameter×height). Adult density is ~70 per vial for the first couple of days of their adult life and thereafter ~2800 individuals in a ~6.4 l cage (19cm×14cm×24cm). We also used a genetically marked population, BL_st_ which was derived from BL_1_ by introducing an autosomal recessive marker – scarlet eye, st (Dasgupta et al. 2016) through a series of six backcrosses. BL_st_ population is maintained under a set of conditions identical to the other BL populations.

### Paternal treatment

Sires and dams were generated from a BL population. The design of the protocol followed to generate the experimental flies is described in Figure 1. To generate the experimental sires and the dams, eggs were collected from a BL population and cultured under standard density (i.e., 70 per 6-8ml food per vial). 100 such vials were set up, of which 65 were used to collect the sires (= sire-vials) and the remaining 35 for dams (dam vials). Dams were collected as virgins and held in single sex vials at a density of 25 per vial with ad lib food until the day of the sire-dam mating (see below). In the sire-vials, all the flies were allowed to eclose. These flies were used to set up three adult density treatments – normal (N: 70 individuals per vial), medium (M: 140 individuals per vial) and high (H: 210 individuals per vial). 10 vials were set up for each of the treatments, using flies that were approximately 1-day old. These vials were left undisturbed for two days, following which males from them were separated and used as sires in the subsequent step. Here and elsewhere throughout the study, all the fly sorting, including collection of virgins, were done under light CO_2_-anaesthesia, unless mentioned otherwise.

**Figure 1:**
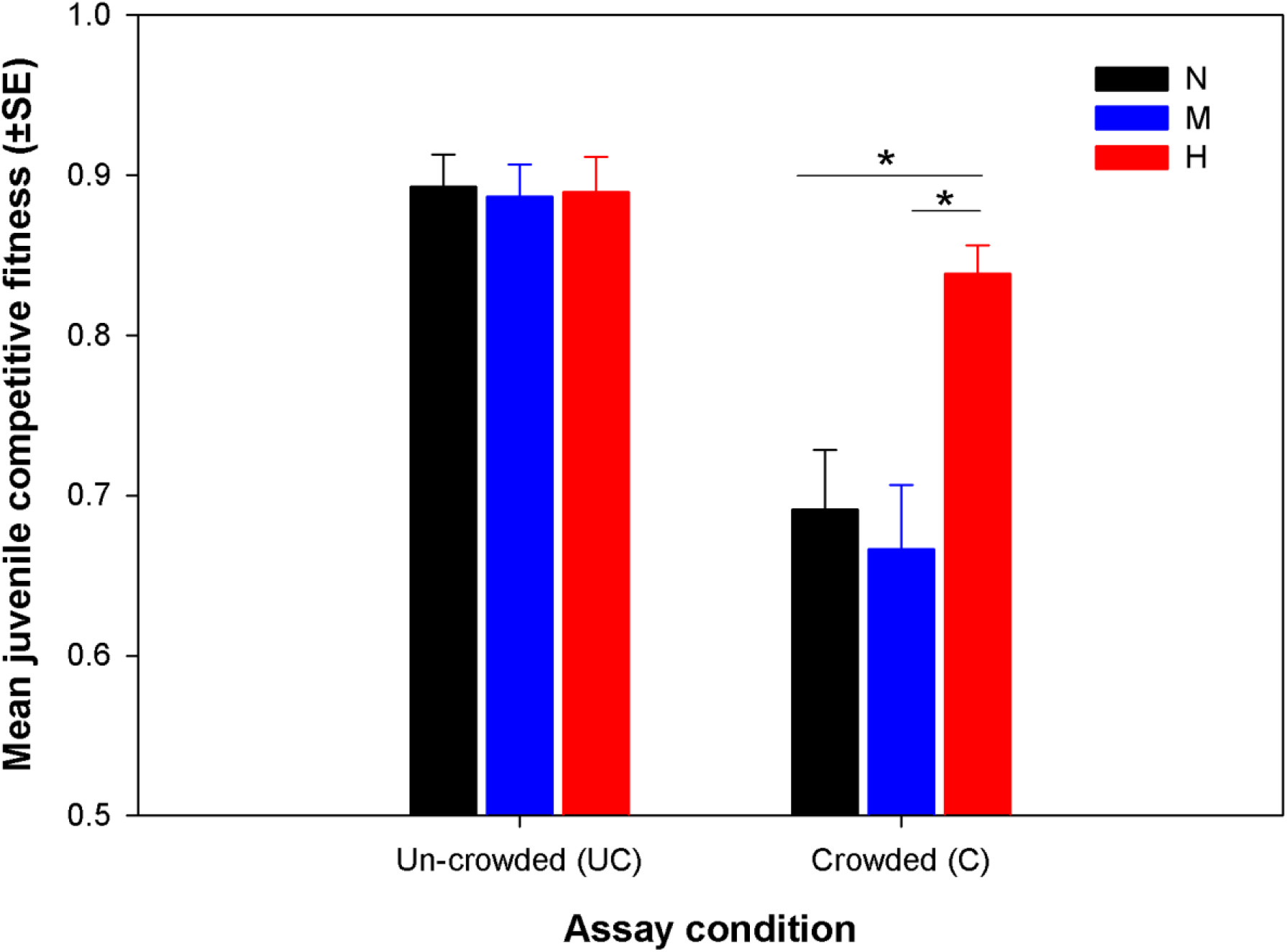
The design of the assay. The schematic diagram shows the design of the entire study, which spanned two generations. Treatment [Normal (N), Medium (M) and High (H) adult densities] was given in the paternal generation. Untreated dams were mated to the treated sires, followed by the collection of eggs from the dams. Assays were done with the eggs and the offspring emerging out of the eggs. Some eggs were subjected to mixed culture (along with competitor eggs) and juvenile competitive fitness vials were set up. Some eggs were cultured as monocultures (without any competitor eggs) – male progeny emerging from these vials were used for further assays, such as, mating ability, mating latency, copulation duration, competitive mating success and courtship frequency.

### Sire-dam mating

Following the 2-day long conditioning, 25 males were randomly isolated from each adult density treatment vials, to be used as sires. They were then combined with dams (see previous section) in fresh food vials (25 sires + 25 dams in a vial) and allowed to interact for 90 minutes, which is sufficient time for a single round of mating. This method of ensuring single round of mating has been previously used (Nandy et al., 2012). In addition, mating was visually observed. Occasionally, in some vials, a small number of females failed to mate within this time. We did not make any attempt to remove them. These un-mated females either mated with an already mated male after a while (late mating) or remained un-mated. Most males secured a single mating, while some very small number (those which mated with the un-mated females mentioned earlier) may have secured more. The number of such late-matings (and hence, male re-mating) was very small, and therefore very unlikely to have any perceivable impact on the subsequent assays. Further, the females in this system usually do not re-mate within such short span (i.e., 90 minutes) unless the first one was a failed mating, which is very rare in our populations. Therefore, by following this protocol, we generated singly inseminated females (average number per vial ~ 25). 10 mating vials were set up per density treatment. After mating, the sires were discarded and the already inseminated dams from all 10 vials of a treatment (i.e., a total of 250 females) were transferred to a 2 litre plastic cage with food smeared with ad-lib quantity of live yeast. Three such cages were thus set up – one for each density treatment. After two days, eggs were collected from these cages to set up the remainder of the experiments. To collect the eggs, a fresh food plate was introduced in the cage. The dams were allowed a short window (2-3 hours) for oviposition. Using a fine brush, eggs were counted on to a fine Agar-strip, which was then transferred to the culture vials (see below). These eggs are hereafter referred to as treatment eggs.

### Measurement of egg-size

To test if the sires influenced the size of the eggs laid by the dams (Pischedda et al., 2010), a subset of these eggs were frozen at -20 °C and their size was measured. For this purpose, eggs were mounted on a glass slide on their dorsal side and photographed using Nikon Stereozoom trinocular microscope (SMZ745T) and the area of the two-dimensional elliptical outline of the eggs were measured in ImageJ, software. This area was taken as a proxy for the size of each egg. A given egg was measured thrice and the average of these three measurements was taken as the unit of analysis. 50 eggs per treatment were measured for this purpose.

### Experiment 1: Juvenile fitness assay

Egg to adult survivorship was taken as a measure of Juvenile fitness. Survivorship of the treatment eggs were measured against a back ground of a common competitor (BL_st_) under two conditions – crowded (C: 150 larvae per 1.5ml food in each vial) and un-crowded (UC: 70 larvae per 6ml food in each vial). During the assay, treatment eggs generated in the previous step were cultured with eggs from common competitors in the ratio 1:4 (C: 30 targets, 120 competitors; UC: 14 targets, 56 competitors). These common competitors were collected from an untreated BL_st_ stock. On completion of development, it was possible to identify the target progeny from the competitor progeny based on eye colour – progeny of the competitors was scarlet eyed whereas the target progeny was red eyed. 10 juvenile competition vials were set up for each of the three treatments (viz., N, M and H) and two assay conditions (i.e., 10 as C and 10 as UC for each treatment). These vials were left undisturbed until adult emergence was complete (12^th^ day post-egg deposition). The adults were sorted based on eye colour and counted. Juvenile fitness score (w) was calculated for each vial following the formula:

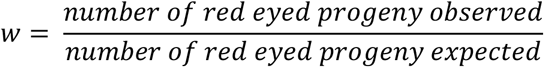

The number of red eyed progeny expected was 14 and 30 for UC and C assay conditions respectively.

### Experiment 2: Assay for behaviour and fitness of the sons

To investigate the effect of the treatment on the male progeny, the treatment eggs were cultured in food vials in the usual density (i.e., 70 per 6 ml food in each vial) and the progeny were allowed to develop. Upon onset of eclosion, males were collected as virgins (< 6 hours post-eclosion). Four assays were run with these males. (a) For each treatment, 50 males were immediately frozen at -20 °C and were later dried at 60 °C for 48 hours and weighed in groups of five using Shimadzu AUW220D to the nearest 0.01mg. (b) A separate set of males were similarly collected and held in groups of 5 per vial for further assays. Ten such vials, for each treatment, were set up and left undisturbed till they were 3days old. These males were then transferred to fresh food vials (hereafter referred to as mating vials) along with five age-matched, virgin females. Mating vials were set up without the use of anaesthesia. The females used in this step came from the same replicate BL population and were generated under their standard maintenance conditions, collected as virgins and held in groups of five per vial with ample food until the day of the experiment. 10 mating vials were set up for each of the three treatments. They were observed (manually, without any video recording) continuously till all the flies finished mating. Every two minutes starting from the time when the females were introduced in these vials, the total number of mating pairs (n_x_, n: number, x: time elapsed in minutes) was noted down at each time point (x = 0, 2, 4, 6…). Mean mating latency (ML, time taken by a virgin pair to start mating) and mean copulation duration (CD, duration for which a pair mated) were calculated following an algorithm mentioned below.

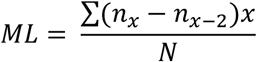

For all values of x, until, *n*_*x*−2_ ≤ *n*_*x*_.

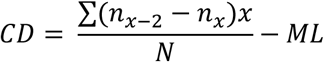

For all values of x, until, *n*_*x*−2_ ≥ *n*_*x*_.

Occasionally, some females did not mate within one-hour long observation. These flies were excluded from the analysis. Similarly, some males also failed to secure mating. In vials having such an unsuccessful male, a mating was recorded much later – when one of the successful males finished its first mating and then initiated a second one with the un-copulated female. Such late copulations were also excluded from the analysis. Mating ability (MA) is measured as the proportion of the sons successfully copulated. MA was calculated for every single vial.

(c) Courtship frequency was quantified for the 3day old (post-eclosion) sons of the three paternal density treatments by setting up similar mating vials as described in the previous section. Ten vials were set up for each treatment. Therefore, a total of 30 vials were observed. After allowing the first mating, the courtship observation was initiated after a gap of approximately half an hour. Vials where all the flies did not mate were removed from the assay. Every 45 minutes, each vial was observed for 30 seconds, during which the total number of courtship bouts (male to female) was noted down. A total of 8 observations were taken. In *Drosophila*, courtship behaviour includes chasing, tapping, courtship dance and song, genital licking and attempted mounting (Bastock & Manning, 1955; Sokolowski, 2010). Any of the above mentioned courtship behaviours, displayed by the five males in each vial was counted as one. The total number of independent male to female courtship displays was counted within the observation window (Nandy et al., 2013). The treatment identities were unknown to the observers to avoid observer bias. (d) Another set of males were similarly collected and held, to be used for quantifying their mating success under competitive condition (CMS, Competitive mating success). This was done by setting up mating vials with five 3day old target males, five competitor males (BL_st_) and five virgin females (BL_st_). Ten such mating vials were set up for each of the three treatments. After allowing a single round of mating for all the females in a mating vial, the females were individually transferred to oviposition test tubes (12mm diameter × 75mm height) with ample food. The females were allowed to oviposit for 18 hours. Following oviposition, the females were discarded and the tubes were retained to allow the progeny to develop and eclose. For each female, the identity of their mate (whether target/competitor) was ascertained by observing the eye colour of the progeny. Progeny sired by target males were red eyed whereas those sired by competitors were scarlet eyed. For a given vial, average CMS of the five target males in the vial was calculated as the proportion of the females mated to target males (i.e., produced red eyed offspring).

### Experimental replications and data analyses

The entire study was carried out in three randomized blocks, using three different BL populations - BL_1_, BL_3_ and BL_5_. The blocks were handled on separate days. Number of replications within each block has been mentioned in the previous sections along with the assay design. Except for the egg size and dry body weight assay, all the experimental replication was done at the level of assay mating vials or juvenile competition vials. All the assays had 10 replicate vials. Vial means were used as the unit of analysis. For egg size assay, size of each egg was used as the unit of analysis. For dry body weight, weight of groups of 5 individuals was used as the unit of analysis. Data were analysed using mixed model Analysis of Variance (ANOVA). Block was treated as random factor, while paternal density treatment and assay density (wherever applicable) were treated as fixed factors. Multiple comparisons were done using Tukey’s HSD. All the analyses were done in Statistica, version 10 (Statsoft, Tulsa, OK, U.S.A.).

## Results

Variation in size of the eggs represents variation in maternal provisioning. The effect of the paternal density treatment on size of the eggs produced by the dams was not significant (Table 1, mean ±SE, μm^2^, N: 80039.1 ±387.5; M: 79967.4 ±415.3; H: 79611.9 ±411.4). The juvenile competitive fitness assay quantified overall egg to adult survival of the target juveniles compared to the same of juveniles from a common background (common competitors). While the data from un-crowded assay condition reflects the baseline survivorship, those from crowded assay condition represents difference in juvenile competitive ability across the three paternal density treatments. Paternal density treatment had a significant effect on Juvenile fitness (Table 1). While there was no significant difference between N and M-treatments, H-treatment had 8.9% higher juvenile fitness compared to that of the N-treatment. This relative advantage of the H-treatment was only evident under larval crowding, i.e., C-assay density (Figure 2), indicating competitive superiority of the H-juveniles. However, the paternal treatment × assay density interaction was marginally non-significant (Table 1).

**Figure 2:**
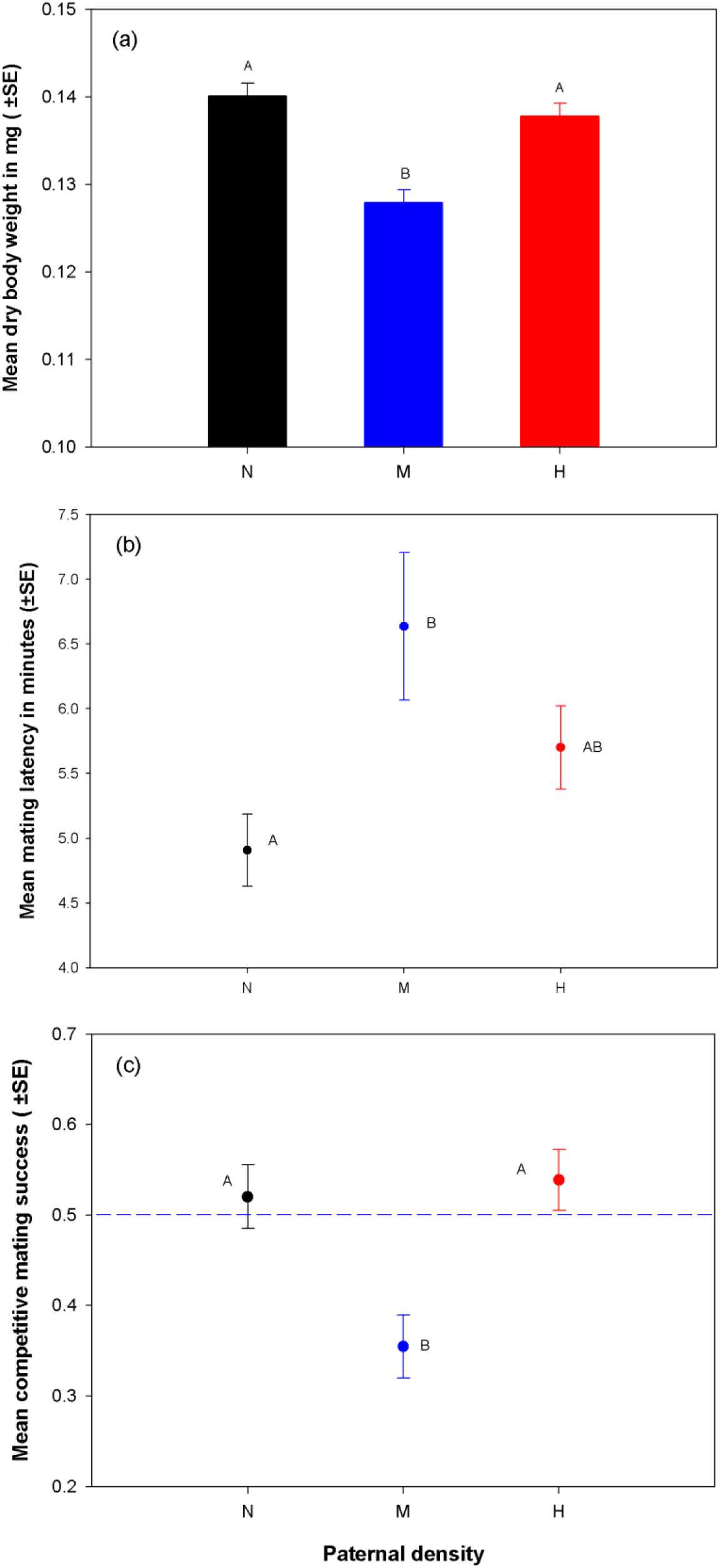
Effect of the paternal density treatment on juvenile competitive fitness, under crowded and un-crowded assay conditions. Target eggs (eggs produced by dams mated to treatment sires) were cultured with competitor eggs (eggs produced by untreated females and males) in juvenile competition vials – under un-crowded and crowded conditions. Proportion of the target progeny successfully emerging as adults is considered as the measure of juvenile competitive fitness. Black, blue and red colour coding represent the progeny of Normal (N), Medium (M) and High (H) density treatment males respectively. The H-progeny were found to have higher juvenile competitive fitness compared to N and M-progeny (represented by *asterisk), only when assayed under crowded condition. The entire experiment was done following a randomized block design and the data were analysed using three factor mixed model ANOVA with paternal treatment and assay condition as fixed factors and block as random factor.

**Table 1:** Summary of results of mixed model ANOVA on the various traits under investigation. Paternal density and assay density (wherever applicable) were considered as fixed factor, while block as random factor. All tests were done considering α=0.05 and significant p-values are mentioned in **bold** face.

| Trait | Effect | SS | DF | MS | DF Den | MS Den | F | p |
| --- | --- | --- | --- | --- | --- | --- | --- | --- |
| Egg size | Paternal density (PD) | $1.75 \times 10^7$ | 2 | $8.75 \times 10^6$ | 4.02 | $3.53 \times 10^6$ | 2.48 | 0.20 |
| | Block | $9.84 \times 10^8$ | 2 | $4.92 \times 10^8$ | 4.03 | $3.54 \times 10^6$ | 139.11 | **<0.01** |
| | PD×Block | $1.41 \times 10^7$ | 4 | $3.53 \times 10^6$ | 438.00 | $2.25 \times 10^7$ | 0.16 | 0.96 |
| Juvenile fitness | Paternal density (PD) | 0.21 | 2 | 0.11 | 4.03 | 0.01 | 9.93 | **0.03** |
|  | Assay density (AD) | 0.97 | 1 | 0.97 | 2.00 | 0.10 | 9.31 | 0.09 |
|  | Block | 0.40 | 2 | 0.20 | 1.58 | 0.09 | 2.15 | 0.35 |
|  | PD×AD | 0.23 | 2 | 0.11 | 4.01 | 0.02 | 5.42 | 0.07 |
|  | PD×Block | 0.04 | 4 | 0.01 | 4.00 | 0.02 | 0.51 | 0.74 |
|  | AD×Block | 0.21 | 2 | 0.10 | 4.00 | 0.02 | 4.96 | 0.08 |
|  | PD × AD × Block | 0.08 | 4 | 0.02 | 148.00 | 0.02 | 1.24 | 0.30 |
| Dry body weight | Paternal density (PD) | $2.67 \times 10^{-7}$ | 2 | $1.34 \times 10^{-7}$ | 4.00 | $6.22 \times 10^{-9}$ | 21.49 | **< 0.01** |
| | Block | $7.78 \times 10^{-7}$ | 2 | $3.89 \times 10^{-7}$ | 4.00 | $6.22 \times 10^{-9}$ | 62.50 | **< 0.01** |
| | PD×Block | $2.49 \times 10^{-8}$ | 4 | $6.22 \times 10^{-9}$ | 80 | $1.27 \times 10^{-8}$ | 0.49 | 0.74 |
| Mating latency | Paternal density (PD) | 41.810 | 2 | 20.90 | 4 | 1.12 | 18.59 | **0.01** |
|  | Block | 51.158 | 2 | 25.58 | 4 | 1.12 | 22.75 | **0.01** |
|  | PD×Block | 4.477 | 4 | 1.12 | 77 | 4.59 | 0.24 | 0.91 |
| Copulation duration | Paternal density (PD) | 14.43 | 2 | 7.21 | 4 | 1.81 | 3.98 | 0.11 |
|  | Block | 80.87 | 2 | 40.44 | 4 | 1.81 | 22.32 | **0.01** |
|  | PD×Block | 7.22 | 4 | 1.81 | 77 | 5.51 | 0.33 | 0.86 |
| Mating ability | Paternal density (PD) | 0.009259 | 2 | 0.00 | 4 | 0.04 | 0.10 | 0.90 |
|  | Block | 0.046678 | 2 | 0.02 | 4 | 0.04 | 0.52 | 0.63 |
|  | PD×Block | 0.179963 | 4 | 0.04 | 77 | 0.02 | 2.64 | **0.04** |
| Courtship frequency | Paternal density (PD) | 13.28 | 2 | 6.64 | 4.03 | 10.79 | 0.62 | 0.58 |
|  | Block | 500.78 | 2 | 250.39 | 4.00 | 10.81 | 23.16 | **0.01** |
|  | PD×Block | 43.24 | 4 | 10.81 | 73.00 | 7.36 | 1.47 | 0.22 |
| Competitive mating success | Paternal density (PD) | 0.64 | 2 | 0.32 | 4.03 | 0.02 | 19.47 | **0.01** |
|  | Block | 0.09 | 2 | 0.05 | 4.06 | 0.02 | 2.86 | 0.17 |
|  | PD×Block | 0.07 | 4 | 0.02 | 75.00 | 0.04 | 0.41 | 0.80 |

We only quantified the effect of the paternal density on male offspring. We found a significant effect of the treatment on dry body weight, ML and CMS (Table 1, Figure 3). Multiple comparisons using Tukey’s HSD indicated that dry body weight of the M-sons were significantly less than that of the N-sons, with M-sons having 8.7% lower mean dry body weight. The difference between the dry body weight of the H and N-sons was not statistically significant. Hence the M-sons were significantly smaller compared to the other two treatments. In the mating assay, though we found some males to fail in acquiring mating, there was no effect of the treatment on mating ability of the sons (MA: mean ±SE, N: 0.91 ±0.04; M: 0.91 ±0.04; H: 0.93 ±0.04). The M-treatment sons showed significantly higher (approximately 35%) ML compared to that showed by the N-treatment sons. While H-treatment also showed 16% higher ML compared to N-treatment, this difference was not significant. Therefore, M-sons took longer to start mating with virgin females indicating females’ reluctance to accept them as mate due to either poor performance in courtship or small size. This relative disadvantage of the M-sons was also evident in terms of their competitive ability in mating competitions. Multiple comparisons on the CMS results indicated that the M-sons had significantly lower CMS compared to H and N-treatments. CMS of the M-sons was approximately 34% less than that of the N-sons. This is however, not due to a reduced courtship performance by the M-sons as we found the effect of the treatment on CF (mean ±SE, N: 6.7 ±0.6; M: 6.8 ±0.6; H: 7.6 ±0.8) to be non-significant. We also did not find any effect of the treatment on CD (mean ±SE, minutes, N: 18.6 ±0.4; M: 17.6 ±0.4; H: 17.9 ±0.4), potentially indicating the lack of the treatment effect on post-copulatory traits of the sons (Table 1).

**Figure 3:**
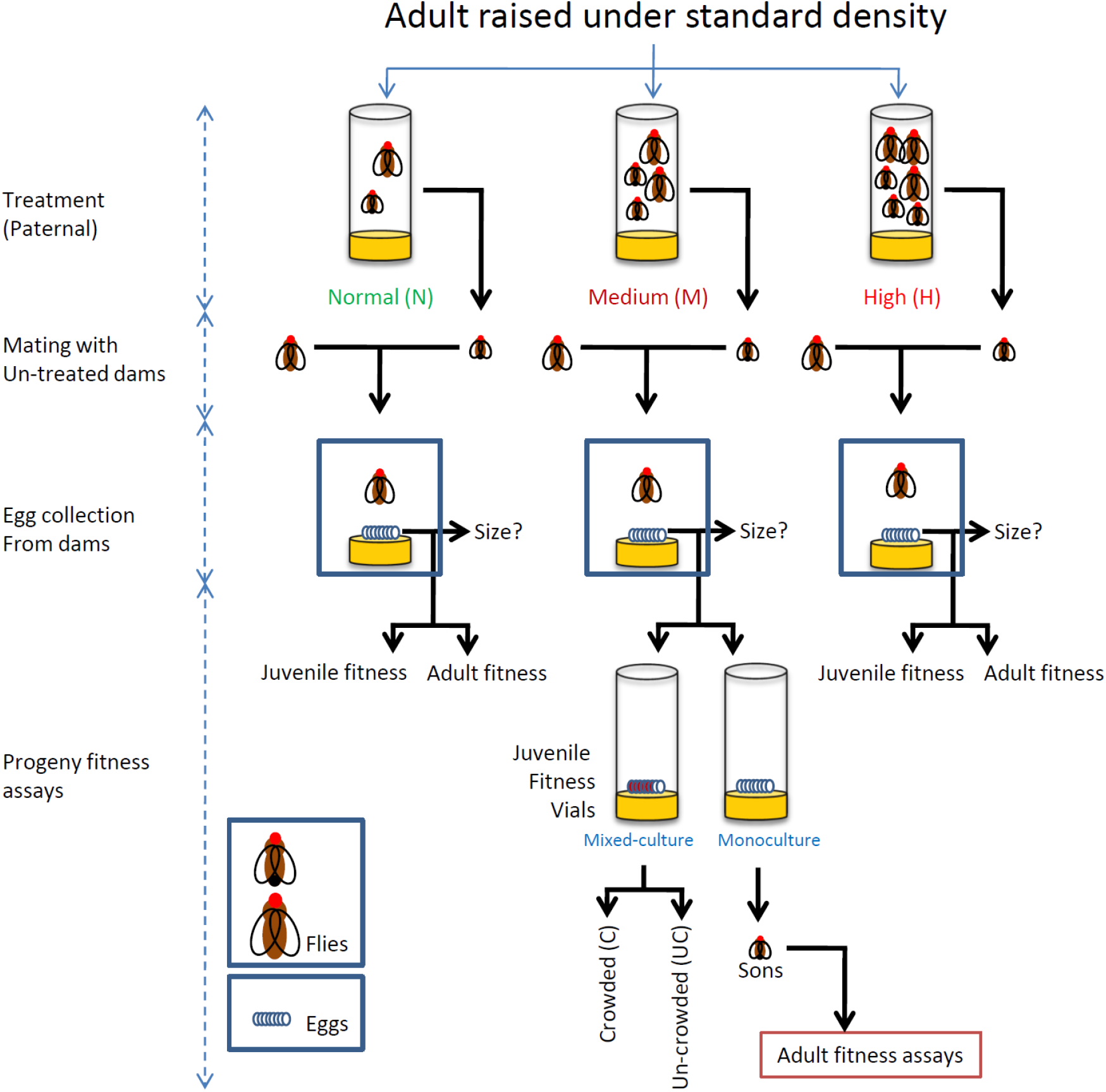
Effect of the paternal density treatment on male offspring. (a) Dry body weight at eclosion: five flies were weighted together to nearest 0.01mg. This was then used as the unit of analysis; (b) Mean mating latency (time taken by a virgin male-female pair to start copulation): mean ML was calculated for five males in a vial following the algorithm given in the Materials & Methods section. This was done for all the mating vials in the assay. These values were then used as the unit of analysis; (c) Competitive mating success (CMS): CMS values were calculated for each vial having five target males as the proportion of females mated to target males in these assay vials. These values were then used as the unit of analysis. The blue broken line indicates the expected value of CMS if there is no mating bias. Black, blue and red colour coding represent data from the progeny of N, M and H males respectively. Treatments not sharing common alphabet were found to be significantly different from each other. The entire experiment was done following a randomized block design and the data were analysed using three factor mixed model ANOVA with paternal treatment and assay condition as fixed factors and block as random factor.

## Discussion

Given that very few studies have shown the effect of paternal environment on offspring fitness components, there were two main objectives of the present study – (a) to assess if paternal exposure to varying population density affected progeny traits; if yes, then (b) to evaluate the adaptive significance of such effect. The results clearly showed that at sufficiently high density, males had an adaptive paternal effect on juvenile competitive fitness. As we did not find any effect of our treatment on size of the eggs produced by the dams, such paternal effect is unlikely to be mediated by variation in provisioning by the females. We further show that at intermediate density, males sire smaller sons which are inferior in acquiring mates. Interestingly, such maladaptive effect of paternal density on offspring adult fitness was not detected at high density.

In holometabolous insects like fruit flies, juvenile (larva and pupa) survival constitutes one of the most important components of fitness (Prasad & Joshi, 2003). In addition, juvenile ecology may also have a major effect on the life-history and fitness components of the adult stage (Heat shock: Khazaeli et al. 1997; cold shock: Singh et al., 2015; Singh & Prasad, 2016; crowding: Joshi & Mueller, 1988; Sarangi et al., 2016; Shenoi et al., 2016). The observed paternal effect on juvenile competitive fitness therefore is extremely consequential. Some relatively recent studies have pointed out that evolving parental ability to optimize offspring fitness related traits can be an adaptation to ecological challenges (Galloway & Etterson, 2007), including crowding (Crean et al., 2013). Given that fruit fly natural ecology regularly involves adult and larval crowding, the observed paternal effect on juvenile competitive fitness can indicate males’ adaptation to crowding. Interestingly, we observed the paternal effect on juvenile competitive fitness, only at the highest density, which may indicate a certain threshold density beyond which such paternal effect starts affecting offspring traits. In addition, when assayed under un-crowded condition the progeny from the three sire treatments do not show any measurable difference in their egg-to-adult survival. This suggests that the juvenile competitive ability rather than baseline juvenile viability was affected by the treatment. Since a number of traits (e.g., feeding rate, waste tolerance, development time etc.) affect juvenile competitive ability in these flies, it will be interesting to find out the trait responsible for better competitive ability of the H-sons in our study.

In a wide range of species including *Drosophila melanogaster*, maternal exposure to high density or poor nutrition has been found to affect offspring fitness components (Prasad & Joshi, 2003; Vijendravarma, 2010; Valtonen, 2012). Such effects can either be beneficial (Mitchell & Read, 2005; Bashey, 2006; Allen et al., 2008; Vijendravarma, 2010; Gorbi et al., 2011) or detrimental (Meylan et al., 2007) depending on the component of the fitness under consideration and the prevailing condition.

As maternal provisioning and other maternal effects play vital roles in offspring survival and performance, such maternal density/nutrition effect is not surprising. However, what is not intuitive is the paternal density to have similar impact on offspring fitness, as our results suggest, given that *Drosophila* males do not pass on any nutrition to the offspring. It is well known in the *Drosophila* literature that even the laboratory populations harbour heritable genetic variation in survival under crowding both as adults and juveniles (see Sarangi et al., 2016 and the references therein for an updated review). Therefore, one possibility is that genetically superior males, which are better at surviving under high density, may produce offspring which are better both as juveniles, explaining at least part of our observations. Though larval competitive ability is known to respond to experimental evolution, indicating heritable genetic variation (Mueller, 1997; Prasad & Joshi, 2003), such heritable variation is very unlikely to have led to the observed treatment effect on juvenile competitive fitness. This is because (a) in our assay, we recorded very little mortality in males during the treatment, indicating negligible hard selection. In addition, we also ensured that there was no soft selection by randomly picking the set of males from the treatment vials to use them as sires. Further, we allowed the sires and the dams to mate only once by allowing them a limited window of time to interact after being put together in mating vials. (b) Even if there was selection in the current experimental design, the selection is likely to be weak (see Materials & Methods section). Such weak selection is unlikely to explain the observed differences in some of the traits (viz., 8.9% increase in juvenile competitive fitness, 35% increase in mating latency), especially within one generation. Alternatively, males may alter maternal provisioning and thereby indirectly affect offspring fitness components (Prasad et al., 2003; Vijendravarma et al., 2010). We, however, did not find any measurable difference in the size of the eggs produced by females mated to the males belonging to the three treatments, making variation in maternal provisioning an unlikely explanation. Therefore, although sire-effect on the quality of the eggs produced by the females cannot be completely ruled out, our results tentatively point at non-genetic paternal effect (Bonduriansky & Day, 2009) as the potential cause behind the observed effect of the treatment. Interestingly, a recent study on *D. melanogaster* has shown paternal effects to have important consequences on the expression of an array of genes in sons (Zajitschek et al., 2017, also see the corresponding correction). In addition, Garcia-Gonzalez & Dowling (2015) reported non-sire effect on daughters’ reproductive output in *D. melanogaster*, possibly caused by the seminal fluid proteins transferred by the males to their mates during copulation (Garcia-Gonzalez & Dowling, 2015).

While we found adaptive paternal effect on juvenile performance, adult performance however, was found to have a significant maladaptive effect of paternal density. Males that experienced intermediate density were found to sire sons which (a) are smaller, (b) take longer time to start mating and (c) have lower mating success. Since we did not find any effect of the treatment on courtship frequency, reduced mating success and increased mating latency was a likely outcome of females’ reluctance to accept relatively smaller males as their mates, a known fitness consequence of reduced size in *Drosophila* males (Partridge et al., 1987; Jagadeeshan et al., 2015). Body size has been reported to be affected by intergenerational paternal effect in another Dipteran – *Telostylinus angusticollis* (Bonduriansky & Head, 2007). Unlike the maladaptive effect found in our study, the paternal effect on body size reported by Bonduriansky & Head (2007), however, was adaptive, especially under certain prevailing conditions. Though *prima facie*, the observed body size reduction appears to be maladaptive, it will be interesting to investigate its fitness consequence under varying adult density. Interestingly, this effect was found only at intermediate density and not in the high density treatment. At this point, it is, however, difficult to suggest any reason for such specific expression of the paternal effect at intermediate density.

As variation in population density and crowding related ecological challenges are common in almost all organisms, including fruit flies, paternal effect of the nature reported here is important to understand. Though paternal ability to optimize offspring traits is likely to be adaptive, especially under fluctuating environment, the results reported here show that paternal effect can be both adaptive and maladaptive. To the best of our knowledge this is the first evidence of the effect of paternal density on juvenile and adult fitness components in *D. melanogaster*. Importantly, our results emphasize the importance of considering paternal effect as a source of variation in fitness related traits. The full impact of such paternal effect in the evolution of life-history traits and the underlying mechanisms are emerging as an important topic of discussion, which is likely to see an increasing attention in years to come.

## Acknowledgements

We are immensely thankful to N. G. Prasad (IISER Mohali) for sharing the fly populations, using which the BL populations were created and also for his critical comments on a previous version of this manuscript. We are also thankful to Syed Zeeshan Ali and Vanika Gupta for their critical comments on a previous version of the manuscript. TV and PD thank Indian Institute of Science Education and Research Berhampur for financial assistance in the form of the Institute Scholarship for Ph.D. programme. The study was supported by a research grant from Department of Science and Technology (INSPIRE Faculty award, Grant no. DST/INSPIRE/04/2013/000520).

## Authors’ contributions

BN, PD and SS conceived the ideas and designed the assays. PD, SS, AAD, TV, and to a lesser extent BN performed the assays and collected the data. Data analysis was primarily done by BN and to some extent, by PD. While BN and PD led the writing of the manuscript, SS and TV provided important assistance and inputs.

## Data intended to be archived in

Dryad

## References

Adler, M.I. & Bonduriansky, R. 2013. Paternal Effects on Offspring Fitness Reflect Father’s Social Environment. Evol. Biol. 40: 288-292.

Allen, R.M., Buckley, Y.M. & Marshall, D.J. 2008. Offspring Size Plasticity in Response to Intraspecific Competition: An Adaptive Maternal Effect across Life-History Stages. Am. Nat. 171: 225-237.

Barnes, A.I. & Siva, M.T. 2000. Density-dependent prophylaxis in the mealworm beetle *Tenebrio molitor* L. (Coleoptera: Tenebrionidae): cuticular melanization is an indicator of investment in immunity. Proc. R. Soc. Lond. B 267: 177-182.

Bashey, F. 2006. Cross-generational environmental effects and the evolution of offspring size in the Trinidadian Guppy *Poecilia reticulate*. Evolution 60: 348-361.

Bastock, M. & Manning, A. 1955 The Courtship of *Drosophila melanogaster*. Behaviour 8: 85-111.

Bonduriansky, R. & Day, T. 2009. Nongenetic inheritance and its evolutionary implications. Annu. Rev. Ecol. Evol. Syst. 40: 103–125.

Bonduriansky, R. & Head, M. 2007. Maternal and paternal condition effects on offspring phenotype in *Telostylinus angusticollis* (Diptera: Neriidae). Int. J. Evol. Biol. 20: 2379-2388.

Christian, J.J. & Lemunyan, C.D. 1958. Adverse effects of crowding on lactation and reproduction of mice and two generations of their progeny. Endocrinology 63: 517-529.

Crean, A.J., Dwyer, J.M. & Marshall, D.J. 2013. Adaptive paternal effects? Experimental evidence that the paternal environment affects offspring performance. Ecology 94: 2575-2582.

Dasgupta, P., Halder, S. & B. Nandy. 2016. Paternal social experience affects male reproductive behaviour in Drosophila melanogaster. J. Genet. 95: 725-727.

Dias, B.G. & Ressler, K.J., 2014. Experimental evidence needed to demonstrate inter-and trans- generational effects of ancestral experiences in mammals. Bioessays 36: 919-923.

Dussourd, D.E., Ubik, K., Harvis, C., Resch, J., Meinwald, J. & Eisner, T. 1988. Biparental defensive endowment of eggs with acquired plant alkaloid in the moth *Utetheisa ornatrix*. Proc. Natl. Acad. Sci. U.S.A. 85: 5992-5996.

Friberg, U., Stewart, A.D. & Rice, W.R. 2012. X- and Y-chromosome linked paternal effects on a life-history trait. Biol. Lett. 8: 71-73.

Galloway, L.F. & Etterson, J.R. 2007. Transgenerational plasticity is adaptive in the wild. Science 318: 1134-1136.

Garcia-Gonzalez F., & Dowling D.K. 2015 Transgenerational effects of sexual interactions and sexual conflict: non-sires boost the fecundity of females in the following generation. Biol. Lett. 11: 20150067.

Gorbi, G., Moroni, F., Sei, S., & Rossi, V. 2011. Anticipatory maternal effects in two different clones of *Daphnia magna* in response to food shortage. J. Limnol. 70: 222-230.

Gwynne, D.T. 1988 Courtship feeding in katydids benefits the mating male’s offspring. Behav. Ecol. Sociobiol. 23: 373-377.

Holbrook, G.L. & Schal, C. 2004. Maternal investment affects offspring phenotypic plasticity in a viviparous cockroach. Proc. Natl. Acad. Sci. U S A. 101: 5595-5597.

Huey, R.B., Wakefield, T., Crill, W.D., & Gilchrist, G. 1995. Within- and between-generation effects of temperature on early fecundity of *Drosophila melanogaster*. Heredity 74: 216-223.

Islam, M.S., Roessingh, P., Simpson, S.J. & McCaffery, A.R. 1994. Parental effects on the behaviour and colouration of nymphs of the desert locust *Schistocerca gregaria*. J. Insect Physiol. 40: 173-181.

Jagadeeshan, S., Shah, U., Chakrabarti, D. & Singh, R.S. 2015. Female Choice or Male Sex Drive? The Advantages of Male Body Size during Mating in *Drosophila Melanogaster*. PLoS ONE 10: e0144672

Joshi, A. & Mueller, L.D. 1988 Evolution of higher feeding rate in Drosophila due to density-dependent natural selection. Evolution, 42: 1090-1093.

Khazaeli, A.A., Tatar, M., Pletcher, S.D. & Curtsinger, J.W. 1997. Heat-induced longevity extension in Drosophila. I. Heat treatment, mortality, and thermotolerance. J. Gerontol. 52a: B48–B52.

Kuijper, B. & Hoyle, R.B. 2015. When to rely on maternal effects and when on phenotypic plasticity? Evolution 69: 950-968.

MacArthur, R.H. & Wilson, E.O. 1967. The theory of island biogeography. Princeton Univ. Press, Princeton, NJ.

Magiafoglou, A. & Hoffmann, A.A. 2003. Cross-generation effects due to cold exposure in *Drosophila serrata*. Funct. Ecol. 17: 664–672.

Mousseau, T. A., & C. W. Fox. 1998. Maternal effects as adaptations. Oxford University Press, Oxford, UK.

Meylan, S., Clobert, J. & Sinervo, B. 2007. Adaptive significance of maternal induction of density- dependent phenotypes. Oikos 116: 650-661.

Mitchell, S.E. & Read, A.F. 2005. Poor maternal environment enhances offspring disease resistance in an invertebrate. Proc. Biol. Sci. 272: 2601–2607.

Mousseau, T.A. & Fox, C.W. 1998. The adaptive significance of maternal effects. Trends in Ecology and Evolution 13: 403-407.

Mueller, L.D. 1997. Theoretical and Empirical Examination of Density-Dependent Selection. Annu. Rev. Ecol. Syst. 28: 269-288.

Mueller, L.D., Guo, P.Z. & Ayala FJ. 1991. Density-dependent natural selection produces trade-offs in life history traits. Science 253: 433-435.

Mueller, L.D. & Sweet, V.F. 1986. Density-dependent natural selection in *Drosophila*: Evolution of pupation height. Evolution 40: 1354-1356.

Nagarajan, A., Natarajan, S.B., Jayaram, M., Joshi, A. 2016. Adaptation to larval crowding in *Drosophila ananassae* and *Drosophila nasuta nasuta*: increased larval competitive ability without increased larval feeding rate. J. Genet. 95: 411-425.

Nandy, B., Gupta, V., Sen, S., Udaykumar, N., Samant, M.A., Ali, S.Z. & Prasad, N.G. 2013. Evolution of mate-harm, longevity and behaviour in male fruit flies subjected to different levels of interlocus conflict. BMC Evol.Biol. 13: 212.

Nandy, B., Joshi, A., Ali, S.Z., Sen, S. & Prasad, N.G. 2012. Degree of adaptive male mate choice is positively correlated with female quality variance. Sci. Rep. 2: 447.

Partridge, L., Ewing, A. & Chandler, A. 1987. Male size and mating success in *Drosophila melanogaster*: the roles of male and female behaviour. Anim. Behav. 35: 555-562.

Pischedda, A., Stewart, A.D., Little, M.K., Rice, W.R. 2010. Male genotype influences female reproductive investment in *Drosophila melanogaster*. Proc. R. Soc. B 278: 2165–2172.

Prasad, N.G. & Joshi, A. 2003. What have two decades of laboratory life-history evolution studies on *Drosophila melanogaster* taught us? J. Genet. 82: 45–76.

Prasad, N.G., Shakarad, M., Rajamani, M. & Joshi, A. 2003. Interaction between the effects of maternal and larval nutritional levels on pre-adult survival in *Drosophila melanogaster*. Evol. Ecol. Res. 5: 903-911.

Reznick, D.N. & Reznick, Y.A. 1993. The influence of fluctuating resources on life-history: Patterns of allocation and plasticity in female Guppies. Ecology 74: 2011-2019.

Rossiter, M.C. 1996. Incidences and consequences of inherited environmental effects. Annu. Rev. Ecol. Syst. 27: 451-476.

Sarangi, M., Nagarajan, A., Dey, S., Bose, J. & Joshi, A. 2016. Evolution of increased larval competitive ability in *Drosophila melanogaster* without increased larval feeding rate. J. Genet. 95: 491-503.

Shenoi, V.N., Banerjee, S.M., Guruswamy, B., Sen, S., Ali, Z.S., & Prasad, N.G. 2016. *Drosophila melanogaster* males evolve increased courtship as a correlated response to larval crowding. Anim. Behav. 120: 183-193.

Shenoi, V.N. & Prasad, N.G. 2016. Local adaptation to developmental density does not lead to higher mating success in *Drosophila melanogaster*. Int.J. Evol. Biol. 29: 407-417.

Singh, K., Kochar, E. & Prasad, N.G. 2015. Egg viability, mating frequency and male mating ability evolve in populations of *Drosophila melanogaster* selected for resistance to cold shock. PLoS ONE 10: e0129992.

Singh, K. & Prasad, N.G. 2016. Evolution of pre- and post-copulatory traits in female *Drosophila melanogaster* as a correlated response to selection for resistance to cold stress. J. Insect Physiol. 91-92: 26-33.

Smedly, S.R. & Eisner, T. 1996. Sodium: A male moth’s gift to its offspring. Proc. Natl. Acad. Sci. U.S.A. 93: 809-813.

Sokolowski, M.B. 2010 Social interactions in “simple” model systems. Neuron 65: 780-794.

Vahed, K. 1998. The function of nuptial feeding in insects: a review of empirical studies. Biol. Rev. 73: 43-78.

Valtonen, T.M., Kangassal, K., Pölkki, M. & Rantala, M.J. 2012. Transgenerational Effects of Parental Larval Diet on Offspring Development Time, Adult Body Size and Pathogen Resistance in *Drosophila melanogaster*. PLoS ONE 7(2): e31611.

Vijendravarma, R.K., Narasimha, S., Kawecki, T.J. 2010. Effects of parental larval diet on egg size and offspring traits in *Drosophila*. Biol. Lett. 6: 238-241.

Watson, M.J.O. & Hoffmann, A.A. 1995. Cross-Generation Effects for Cold Resistance in Tropical Populations of *Drosophila melanogaster* and *Drosophila simulans*. Aust. J. Zool. 43: 51–58.

William, R.R., Stewart, A.D., Morrow, E.H., Linder, J.E., Orteiza, N. & Byrne, P.G. 2006. Assessing sexual conflict in the *Drosophila melanogaster* laboratory model system. Philos. Trans. R. Soc. Lond. B. Biol. Sci. 361: 287-299.

Zeh, D.W. & Smith, R.L. 1995. Paternal investment by terrestrial arthropods. Am. Zool. 25: 785-805.

Zajitschek, F., Zajitschek, S. & Manier, M., 2017. High-protein paternal diet confers an advantage to sons in sperm competition. Biol. lett. 13: 20160914.

Yehuda, R., Bierer, L.M., Schmeidler, J., Aferiat, D.H., Breslau, I. & Dolan, S., 2000. Low cortisol and risk for PTSD in adult offspring of holocaust survivors. Am. J. Psychiatry 157: 1252-1259.

